# D-type K+ current controls the function of electrically coupled neurons in a species-specific fashion

**DOI:** 10.1101/2023.01.06.523023

**Authors:** Antonella Dapino, Federico Davoine, Sebastian Curti

## Abstract

Electrical synapses supported by gap junctions, are known to form networks of electrically coupled neurons in many regions of the mammalian brain, where they play relevant functional roles. Yet, how electrical coupling support sophisticated network operations, and the contribution of the intrinsic electrophysiological properties of neurons to these operations, remains incompletely understood. Here, comparative analysis of electrically coupled mesencephalic trigeminal (MesV) neurons, uncovered remarkable difference in the operation of these networks in highly related species. While spiking of MesV neurons might support the recruitment of coupled cells in rats, this rarely occurs in mice. Using whole-cell recordings, we determined that the higher efficacy in postsynaptic recruitment in rat’s MesV neurons does not result from coupling strength of larger magnitude, but instead from the higher excitability of coupled neurons. Consistently, MesV neurons from rats present a lower threshold current for activation, more hyperpolarized firing level as well as a higher ability to generate repetitive discharges, in comparison to their counterparts from mice. This difference in neuronal excitability results from a significantly higher magnitude of the D-type K+ current (ID) in MesV neurons from mice, indicating that the expression level of this current gates the recruitment of postsynaptic coupled neurons. Since MesV neurons are primary afferents critically involved in the organization of orofacial behaviors, such mechanism might support lateral excitation, by which activation of single neurons at the periphery can spread to coupled partners. Thus, by amplifying sensory inputs, lateral excitation may significantly contribute to information processing and organization of motor outputs.

## INTRODUCTION

Electrical synaptic transmission is a modality of communication mediated by gap junctions, which represent areas of close apposition between the plasmatic membrane of neurons characterized by the presence of clusters of intercellular channels (Bennett, 1997). These junctions establish pathways of low resistance for the flow of ionic currents, supporting the bidirectional and fast communication of both, depolarizing and hyperpolarizing signals between neurons (Connors & Long, 2004; Bennett & Zukin, 2004; Pereda *et al*., 2013). These characteristics allow electrical synapses to contribute to relevant functional operations by neural circuits in many brain areas. Among these operations, lateral excitation results from the ability of electrical coupling to share excitation within neural circuits, creating functional compartments for signal pooling. By such mechanism, the activity in some cells promotes the activation of neighboring coupled ones, thus, operating as a positive feedback mechanism. Although this operation might degrade spatial specificity within neural circuits, lateral excitation between primary afferents tuned to qualitatively similar stimuli, acts to boost sensory responses involved in the organization of motor outputs as was shown in invertebrates and lower vertebrates (El Manira *et al*., 1993; Pereda *et al*., 1995; Herberholz *et al*., 2002; Antonsen *et al*., 2005; Curti *et al*., 2022). In mammals, this mechanism has also been implicated in the enhancement of excitability in circuits of electrically coupled neurons of the olfactory bulb (Christie & Westbrook, 2006), the cerebellum (Vervaeke *et al*., 2012) and the dorsal cochlear nucleus (Apostolides & Trussell, 2013, 2014). Moreover, in the retina, lateral excitation provides a mechanism for precise detection of the spatial location of the stimulus, disregarding its velocity (Trenholm *et al*., 2013).

Mesencephalic trigeminal trigeminal (MesV) neurons are a special class of primary afferents, whose cell bodies, instead of being located in peripheral ganglia, are distributed in the brainstem (Morquette *et al*., 2012). The peripheral branches of these pseudounipolar neurons innervate the spindles of jaw-closing muscles and mechanoreceptors of periodontal ligaments, whereas the central processes supply sensory input to neurons of the trigeminal motor nucleus, the rostral parvocellular reticular formation, and the nucleus supratrigeminalis (Dessem & Taylor, 1989; Liem *et al*., 1991). On the other hand, MesV neurons receive synaptic input at their somata from the hypothalamus and various brainstem structures (Lazarov, 2002; Verdier *et al*., 2004), supporting the notion that these afferents not only relay peripheral information, but are also integral part of the central circuit involved in the generation of masticatory patterns (Morquette *et al*., 2012). Previous work has shown that MesV neurons are electrically coupled by means of large connexin36 (Cx36) containing somato-somatic contacts, and rather than extensive as observed in most structures of the mammalian brain (Connors & Long, 2004; Bennett & Zukin, 2004), coupling in this nucleus is restricted to pairs or small clusters of neurons (Baker & Llinás, 1971; Curti *et al*., 2012). Moreover, the dynamic interaction of electrical coupling with their intrinsic electrophysiological properties, supports the strong synchronization of pairs of MesV neurons and allows them to operate as coincidence detectors (Curti *et al*., 2012; Davoine & Curti, 2019).

A previous study based on dye transfer experiments and electrophysiological recordings, showed that the incidence of coupling in the MesV nucleus is about three times higher in mice compared to rats (Curti *et al*., 2012). This contrasting circuital organization in homologous circuits subserving the same function in highly related species is surprising, raising the possibility that electrical contacts also present functional differences between both species. Particularly, we focused on the ability of presynaptic spikes to recruit postsynaptic neurons, as it represents a critical aspect for operations like synchronization and lateral excitation. Strikingly, here we show that despite similar coupling strength, postsynaptic recruitment in rats is dramatically more efficient than in mice. Such difference does not result from dissimilarities of the gap junctions themselves, but instead from the properties of the non-synaptic membrane of coupled neurons between species. More specifically, combining electrophysiological and pharmacological evidence with a comparative approach, we show that the observed interspecific difference results from a significantly higher functional expression of the D-type K+ current in MesV neurons from mice. These results identify a role for the subthreshold K+ currents in determining the efficacy of postsynaptic recruitment at electrical synapses, and hence the mode of operation of networks of coupled neurons. This emphasizes the role of voltage-dependent membrane conductances in circuits of electrically coupled neurons, and its possible contribution to early stages of sensory processing by primary afferents involved in the organization of relevant behaviors.

## METHODS

### Ethical approval

Sprague-Dawley rats (age: P7 – P16) and C57BL mice of either sex (age: P12 – P18) were obtained from the University animal facility accredited by the local authorities (CHEA: Comisión Honoraria de Experimentación Animal of Universidad de la República, Uruguay). All animal care and experimental procedures were performed in accordance with national guidelines and laws, with minimization of the numbers of animals used. Protocols were approved CNEA (Comisión Nacional de Experimentación Animal) and the School of Medicine CEUA (Comisión de Ética en el Uso de Animales, protocols number: 070153-000128-20 and 070153-000396-17).

### Slice preparation and electrophysiology

Animals were decapitated without anesthesia, and brains were quickly removed. Transverse brainstem slices (180-250 μm thick) were prepared using a vibratome (Leica VT 1000s or DSK DTK-1000) in cold sucrose solution containing 248 mM sucrose, 26 mM NaHCO3, 10 mM glucose, 2 mM MgSO4, 2.69 mM KCl, 1.25 mM KH2PO4 and 1 mM CaCl2 for rats, and 213 mM sucrose, 26 mM NaHCO3, 10 mM glucose, 2 mM MgSO4, 2.69 mM KCl, 1.25 mM NaH2PO4, 1 mM CaCl2, 0.35 mM ascorbic acid and 0.3 mM pyruvic acid for mice. In both cases solutions were bubbled with 95% O2 and 5% CO2 (pH ∼ 7.4). The slices were then transferred to an incubation chamber filled with physiological solution containing 124 mM NaCl, 2.69 mM KCl, 1.25 mM KH2PO4, 26 mM NaHCO3, 10 mM Glucose, 2 mM CaCl2, and 2 mM MgSO4 bubbled with 95% O2 and 5% CO2 (pH ∼ 7.4) at 34°C for 30 min. Afterwards, slices were kept at room temperature in the physiological solution until they were transferred into the recording chamber. The recording chamber, mounted on an upright microscope stage (Nikon Eclipse E600), was continuously perfused with physiological solution (1 – 1.5 ml/min) at room temperature. Whole cell patch recordings were performed under visual control using infrared differential interference contrast optics (IR-DIC). MesV neurons were identified on the basis of their location, large spherical somata, and characteristic electrophysiological properties in response to both depolarizing and hyperpolarizing current pulses (Curti *et al*., 2012). Recording pipettes pulled from borosilicate glass (4 – 8 MΩ) were filled with intracellular solution containing 148 mM K-gluconate, 3 mM MgCl2, 0.2 mM EGTA, 4 mM Na2-ATP, 0.3 mM Na-GTP and 10 mM HEPES (pH ∼ 7.2). The seal resistance between the electrode tip and the cell membrane was higher than 1 GΩ and pipette capacitance was compensated before breaking the seal. Simultaneous recordings from pairs of MesV neurons whose cell bodies lie in close apposition were made using a Multiclamp 700B amplifier (Molecular Devices, Sunnyvale, CA). Under current clamp configuration, the voltage drop across the microelectrode resistance was eliminated by means of the bridge balance control of the amplifier. In voltage clamp configuration, the membrane capacitance and series resistance were compensated (80%) and continuously monitored. Only cells displaying resting membrane potential more negative than -50 mV, or spike amplitude above 70 mV were included in this study. Recordings were low-pass-filtered at 5 kHz and acquired by means of an analog to digital converter connected to a computer, sampled at 10 - 40 KHz depending on the experiment.

### Calculation of coupling coefficient (CC)

During simultaneous whole cell recordings of pairs of coupled MesV neurons in current clamp, a series of hyperpolarizing current pulses of different amplitudes (−50 to -450 pA) were applied to one cell, whereas the voltage changes were recorded in the same injected neuron (presynaptic, Vpre) and in the coupled one (postsynaptic, Vpost). Plots of Vpost as a function of Vpre were constructed and CC was estimated from the slope of linear regressions. For each pair the CC was estimated in both directions.

### Calculation of the input resistance (Rin)

During simultaneous whole cell recordings of pairs of coupled MesV neurons in current clamp, a series of hyperpolarizing current pulses of different amplitudes (−50 to -450 pA) were alternatively applied to one or the other cell, whereas the voltage changes were recorded in the same injected neuron (Vm). Plots of Vm as a function of the intensity of injected current were constructed and the Rin was estimated from the slope of linear regressions. This value represents the equivalent resistance of two parallel branches, one corresponding to the non-junctional resistance of the injected neuron and the other to the gap junction plus the non-junctional resistance of the coupled neuron (Bennett, 1966).

### Estimation of gap junction conductance

From current clamp recordings, the conductance of electrical contacts (Gj) was estimated as the reciprocal of the resistance (Rj) calculated according to the following equation (Bennett, 1966):

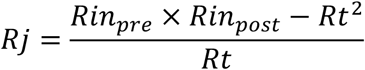

where Rin_pre_ and Rin_post_ are the input resistance of the pre- and postsynaptic cells respectively and Rt is the transfer resistance defined as the voltage response amplitude in the coupled postsynaptic cell divided by the current amplitude intensity injected in the presynaptic cell. Conductance values estimated by this method are reported as directions (two directions per coupled pair).

### Frequency-transfer analysis

The transfer properties between pairs of electrically coupled MesV neurons was determined by injecting frequency modulated (2– 600 Hz or 0.5 to 100 Hz) sine waves of current into one of the coupled cells (ZAP protocol), while recording the resulting membrane voltage deflections in both cells (see Fig. 4A, inset). Peak-to-peak intensity (50 – 300 pA) was adjusted to induce subthreshold voltage deflections. Magnitude of fast Fourier transform (FFT) was calculated for presynaptic and postsynaptic membrane responses, and the frequency-transfer property was determined as the ratio of the postsynaptic FFT magnitude over the presynaptic FFT magnitude. The population frequency-transfer function for both species was determined by averaging single transfers in each recorded direction. Average transfer functions were low-pass filtered applying a smoothing algorithm, and expressed in decibels (dB). The apparent cutoff frequency was determined as the intersection of the slope of the attenuation observed at high frequencies and a horizontal line representing the value observed in DC.

### Assessment of MesV neurons excitability

During current clamp recordings, series of depolarizing current pulses of 200 ms in duration were applied, whose intensities ranged from 50 to 600 pA, in steps of 50 pA. From these recordings, curves of the number of spikes versus current intensity were constructed and the slope of these relations were determined by linear regression analysis forcing the fitting through the origin. This slope reflects the ability of the neuron to produce repetitive discharges as well as the threshold current, representing a valuable indicator of neuronal excitability (Davoine & Curti, 2019).

### Recording K+ currents (IA and ID)

In order to reduce current intensity through voltage-gated Na+ channels and HCN channels, NaCl was substituted by Choline-Cl and TTX (0.25 μM) and CsCl (2.5 mM) were added to the physiological solution. In this condition (control), a series of step-like voltage commands of 500 ms in duration, from 0 to 70 mV in steps of 5 mV starting from a holding potential of -70 mV, was simultaneously applied to both recorded neurons. These voltage commands were repeated after the addition of 4-aminopyridine (4-AP, 30 μM) and of 4-AP (1 mM). The ID was isolated subtracting current traces obtained after the addition of 4-AP (30 μM) from those obtained in control, whereas the IA was isolated subtracting current traces obtained after the addition of 4-AP (1 mM) from those obtained in the presence of 4-AP (30 μM) (Storm, 1988; Mitterdorfer & Bean, 2002). From current traces obtained following this procedure, peak values within the first 50 ms of voltage commands were determined, and transformed to conductance values dividing by the corresponding driving force for K+, in order to construct activation curves for the ID and the IA. For the construction of inactivation curves, conductance values were determined from maximum currents evoked by 10 ms voltage steps to -30 mV, preceded by voltage commands of 500 ms in duration, from -70 mV to 0 mV in steps of 5 mV. Activation and inactivation curves were fitted to a Boltzmann equation of the form:

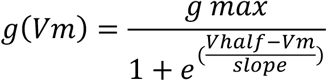

From these fits, maximum conductance (gmax), half activation voltage (Vhalf) and slope at Vhalf were obtained. To avoid bias by difference in cell size, gmax was also reported normalized by the cell’s membrane capacitance. The IA and ID half activation voltages were significantly different both in rats (IA Vhalf: -27.9±4.4 mV [SD], n=17; ID Vhalf: - 39.2±3.8 mV [SD], n=20; P=1.42 × 10^−9^), and mice (IA Vhalf: -28.9±5.2 mV [SD], n=14; ID Vhalf: -36.3±4.2 mV [SD], n=13; P=0.0004), indicating that these currents were successfully separated following the procedure described above.

### Recording of the persistent Na+ current (INap)

To isolate the INap, currents through K+ channels and HCN channels were reduced by adding a combination of blockers to the extracellular solution (10 mM TEA-Cl, 1 mM 4-AP and 5 mM CsCl). In this condition (control), series of step-like voltage commands of 500 ms in duration, from 0 to 70 mV in steps of 5 mV, starting from a holding potential of -70 mV, were simultaneously applied to both recorded neurons. These voltage commands were repeated after addition of TTX (0.5 μM). Current traces obtained in TTX were subtracted from those obtained in control, and current intensity was measured after 100 ms of the initiation of the voltage commands. To construct steady-state activation curves, current values were transformed to conductance dividing by the calculated driving force for Na+, and the parameters characterizing this process (gmax, Vhalf and slope at Vhalf) were determined following the same procedure described for the study of K+ currents.

As no obvious difference in cellular excitability was observed between coupled and uncoupled neurons from both species, Na+ and K+ membrane currents were recorded in coupled and uncoupled neurons, and data was pooled. However, in the case of coupled neurons, voltage commands were always simultaneously delivered to both neurons in order to improve space clamp.

### Data analysis and statistics

Data was analyzed using the following software: Axograph X, Igor Pro7 (Wave Metrics) and Python scientific development environment Spyder (libraries: Numpy, Scipy, Axographio, Stfio, Pandas and Matplotlib). Results were expressed as average value ± standard deviation (SD). Significance of quantitative data was determined by using Student’s t-test of Igor Pro7 (Wave Metrics).

## RESULTS

Previous work had shown that the incidence of electrical coupling among MesV neurons from rats and mice is dramatically different. Both electrophysiological and tracer coupling experiments revealed that coupling in rats is about 21-23%, whereas in mice it is considerably more prevalent being about 60-63% (Curti *et al*., 2012). Such findings suggest contrasting principles of network organization in homolog rodent circuits involved in the organization of orofacial behaviors. To determine if such rat-mouse difference in network organization is accompanied by functional dissimilarities, under identical recording conditions, the properties of electrical synaptic transmission between MesV neurons from these species were systematically compared. Most specifically, the ability of presynaptic spikes to recruit postsynaptic neurons was taken as a readout of functional specializations on the basis of two considerations. First, action potentials triggered in response to sensory stimuli at the periphery most probably constitute the main signal source for coupling between these primary afferents. Second, relevant functional operations of electrical coupling, like synchronization and lateral excitation, rely on the ability of presynaptic neurons to recruit electrically coupled neighbors (Connors & Long, 2004; Pereda *et al*., 2013; Connors, 2017; Curti *et al*., 2022). Thus, the efficacy of postsynaptic recruitment was assessed in pairs of electrically coupled MesV neurons from rats and mice. For this, just-suprathreshold depolarizing current pulses were alternatively injected into each neuron of an electrically coupled pair, whereas the membrane potential was simultaneously monitored in both cells. These experiments revealed that rat’s MesV neurons drives spiking of its postsynaptic coupled neurons in half of the tested directions (35 out of 70 directions, two directions per pair) (Fig. 1A-C), consistent with previous work showing strong spiking synchronization in coupled pairs (Curti *et al*., 2012). In striking contrast, the recruitment of postsynaptic neurons by presynaptic spikes occurs only in 2.4% of tested directions (3 out of 125) in mice (Fig. 1D-F), despite coupling strength of these two populations were matched (see below).

**Figure 1.**
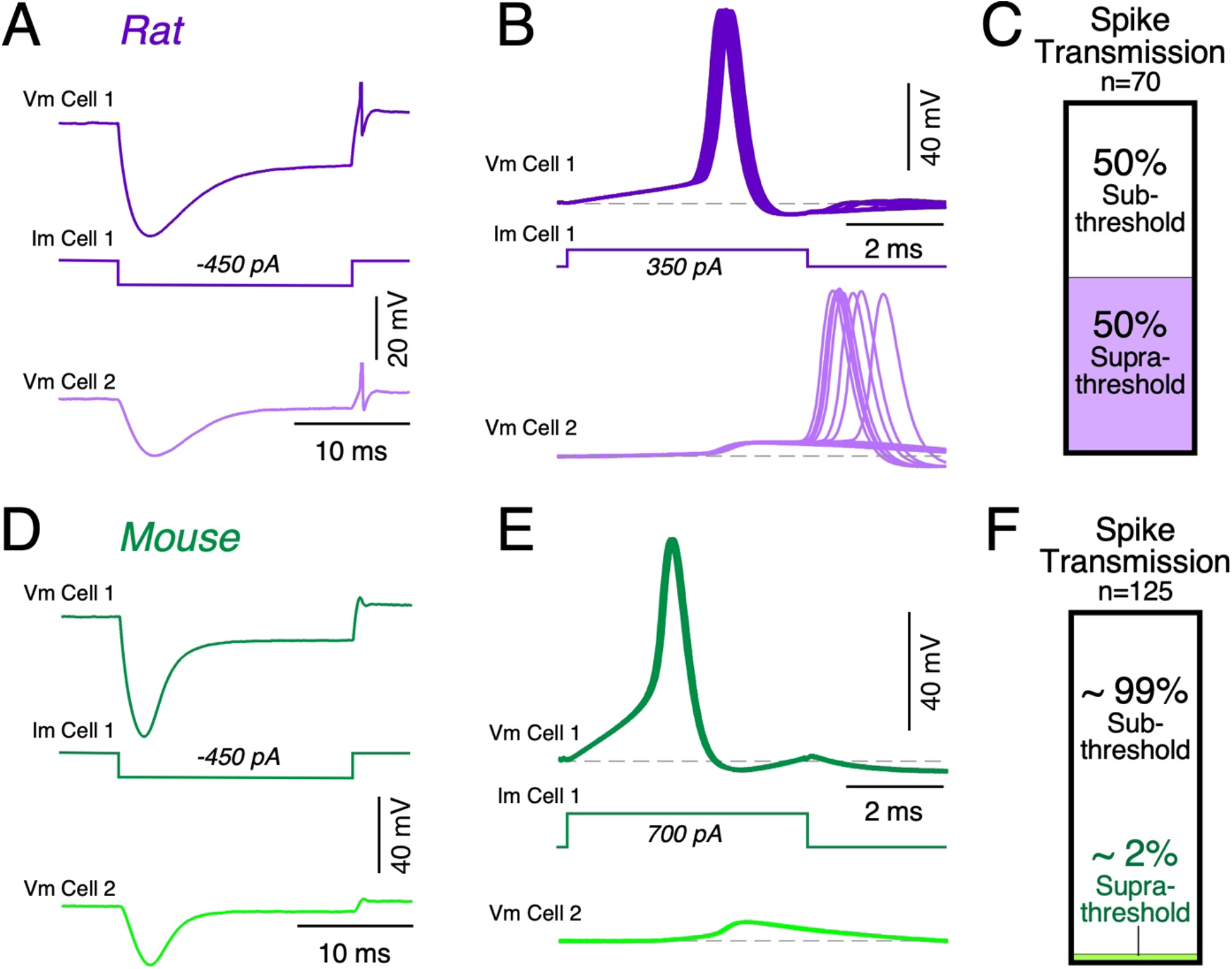
Postsynaptic recruitment between electrically coupled MesV neurons is considerably more efficient in rats compared to mice. *A* and *D*: Paired recordings from electrically coupled MesV neurons during the injection of a hyperpolarizing current pulse (Im Cell 1) showing single traces of the voltage membrane response in the injected (Vm Cell 1) and coupled (Vm Cell 2) neurons, in rat and mouse respectively. For the same pairs depicted in *A* and *D*, postsynaptic recruitment was assessed by activating the presynaptic neuron with a short depolarizing current pulse (Vm Cell 1) in rat (*B*) and mouse (*E*). 50 – 100 single traces are shown superimposed to estimate the firing probability of the postsynaptic neuron in each species. *C* and *F*: Bar graph illustrating the efficacy of postsynaptic recruitment for the population of coupled pairs, as the fraction of tested directions in which an action potential in one neuron induced firing in the other one (Suprathreshold) or not (Subthreshold), in rats and mice (rat: 70 directions, 35 pairs, 34 animals; mouse: 125 directions, 64 pairs, 55 animals).

The higher efficacy of presynaptic spikes to drive spiking in postsynaptic coupled MesV neurons from rats compared to mice, might result from difference in coupling strength between these two species. In order to avoid any bias due to such difference, the populations of coupled pairs from rats and mice were matched in terms of their coupling coefficients (CC). Accordingly, the CC determined by a series of hyperpolarizing current pulses (Fig. 2A and B, see Methods) showed no statistical difference between these two species, averaging 0.44±0.18 [SD] (n=70) and 0.44±0.11 [SD] (n=128), for rats and mice respectively (P=0.981, unpaired, two-tailed *t*-test). Histograms in Figure 2C show that CC from mice display a roughly symmetric distribution, while that from rats is slightly skewed to the right, although this minor difference most probably cannot explain the dramatic dissimilarity in postsynaptic recruitment between these two populations. Consistently, neither of the determinants of the CC, the junctional conductance (Gj) and the input resistance (Rin) of the postsynaptic neuron (Bennett, 1966 p.196; Curti & O’Brien, 2016), displayed any statistical difference. The Rin averaged 92.3±25.3 MΩ [SD] (n=70) and 89.9±27.1 MΩ [SD] (n=128) in rats and mice respectively (P=0.538, unpaired, two-tailed *t*-test) (Fig. 2D), whereas the junctional conductance (Gj) estimated from current clamp recordings (Methods), averaged 7.54±5.64 nS [SD] (n=70) in rats, and 6.64±2.56 nS [SD] (n=128) in mice (P=0.210, unpaired, two-tailed *t*-test) (Fig. 2E). Also, in the age range employed in this study, neither CC nor Rin showed any correlation with age as indicated by linear regression analysis (rat: CC vs age slope=0.011, R^2^=0.022; mouse: CC vs age slope=0.0036, R^2^=0.0013; rat: Rin vs age slope=0.76, R^2^=0.0047; mouse: Rin vs age slope=-1.8, R^2^=0.0057). Thus, disparity in the efficacy postsynaptic recruitment in rats versus mice, cannot be explained by difference in the coupling strength between these species.

**Figure 2.**
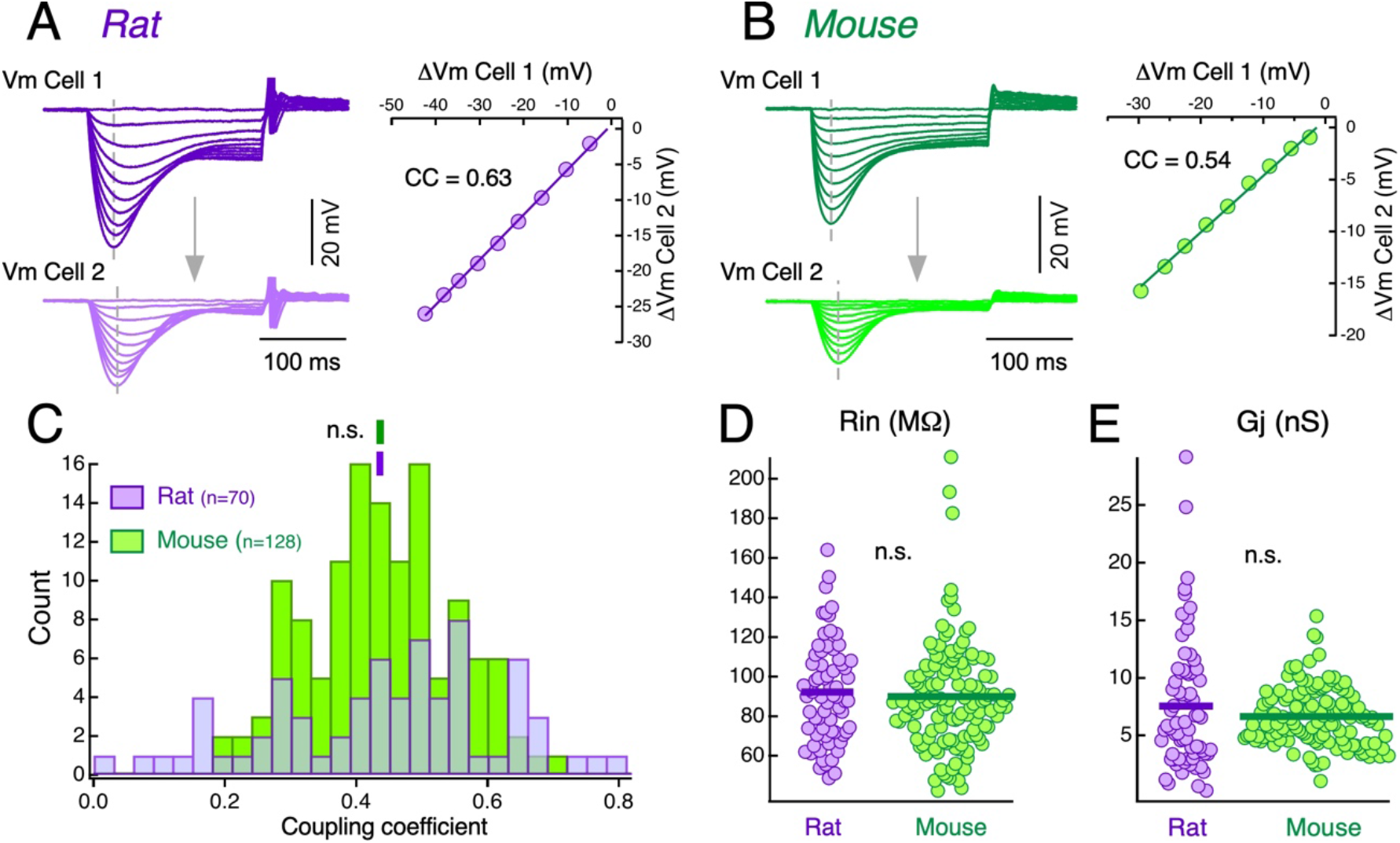
Characterization of the coupling strength and its determinants. The injection of a series of hyperpolarizing current pulses of increasing intensity into one cell produces corresponding voltage responses in the same cell (Vm Cell 1) and in the coupled cell (Vm Cell 2), in rat (*A*, left) and mouse (*B*, left). From these recordings, the coupling coefficient was estimated by plotting the amplitude of membrane voltage changes (measured at the peak of hyperpolarizing responses, vertical dashed lines) in the postsynaptic cell (Vm Cell 2, ordinates) as a function of membrane voltage changes in the presynaptic cell (Vm Cell 1, abscissas). Each data set was fitted with a straight-line function, and the slope value representing the coupling coefficients (CC) are indicated (*A* and *B*, right). *C*: Histogram showing the distribution of CC calculated for the population of recorded directions in rats (purple) and mice (green). Vertical bars above histograms indicate the population average for each data set (rat: 0.44±0.18 [SD], n=70; mouse: 0.44±0.11 [SD], n=128; P=0.981, unpaired, two-tailed *t*-test). *D*: Input resistance (Rin) measured in the population of recorded neurons in rats and mice (rat: 92.3±25.3 MΩ [SD], n=70; mouse: 89.9±27.1 MΩ [SD], n=128; P=0.538, unpaired, two-tailed *t*-test). *E*: Junctional conductance (Gj) values estimated in each assessed direction in rats and mice (rat: 7.54±5.64 nS [SD], n=70; mouse: 6.64±2.56 nS [SD], n=128; P=0.210, unpaired, two-tailed *t*-test). Horizontal bars in *D* and *E* represent population averages.

Although the CC estimated at the peak of hyperpolarizing voltage responses to current pulses is informative about the coupling strength in the passive regime, it might not faithfully reflect the efficacy of spike transmission. On one hand, transmission at electrical synapses typically behaves as a low-pass filter, meaning that high frequency signals, like action potentials, are considerably more attenuated in comparison to signals with a lower frequency content. This implies that besides the Gj and the Rin of the postsynaptic cell, the waveform of the presynaptic signal (i.e., its frequency content) also represent a critical determinant of the coupling strength (Bennett, 1966; Connors & Long, 2004; Alcamí & Pereda, 2019; Curti *et al*., 2022). On the other hand, active subthreshold mechanisms can crucially shape postsynaptic responses. For example, it has been well established that the persistent Na+ current (INap) is able to significantly increase the efficacy of spike transmission at electrical contacts (Mann-Metzer & Yarom, 1999; Curti & Pereda, 2004; Dugué *et al*., 2009; Curti *et al*., 2012).

Thus, in order to assess the efficacy of spike transmission in rats and mice, spike characteristics were determined firstly and their impact on the postsynaptic cell was evaluated thereafter. Figures 3A and B display representative spikes (top) recorded in MesV neurons from rats and mice respectively, with their corresponding time derivatives (bottom), illustrating the method employed to measure several spike parameters. While spike duration does not show statistical difference between rats and mice, averaging 0.48±0.14 ms [SD] (n=69) and 0.50±0.15 ms [SD] (n=117) respectively (P=0.316, unpaired, two-tailed *t*-test) (Fig. 3C), spike amplitude is significantly larger in rats (88.7±11.9 mV [SD], n=70) compared to mice (68.2±12.3 mV [SD], n=118) (P=1.29 × 10^−21^, unpaired, two-tailed *t*-test) (Fig. 3D). Moreover, spike after-hyperpolarization (AHP) is also of larger amplitude in rats compared to mice (rat: -5.8±2.5 mV [SD], n=69; mouse: - 4.9±2.1 mV [SD], n=118; P=0.019, unpaired, two-tailed *t*-test) (Fig. 3E), most probably due to the stronger activation of repolarizing mechanisms during rat action potentials, whose peak levels attain more positive values compared to mice. Thus, although similar in duration, MesV neurons’ spikes from rats are on average ∼20 mV bigger in amplitude than those from mice. Correspondingly, the maximum value of the spike time derivative is significantly higher in rats versus mice (rat: 310.8±96.6 mV/ms [SD], n=66; mouse: 183.5±89.6 mV/ms [SD], n=118; P=9.3 × 10^−15^, unpaired, two-tailed *t*-test), indicating that spikes from rat MesV neurons present a faster time course. Consistently, spike phase plots (first time derivative of membrane voltage as a function of the membrane voltage) show that trajectories from mice lie almost completely within those of rats’ MesV neurons (Fig. 3F). Also, this analysis revealed that the firing level, defined as the value of membrane voltage at which the rate of change reaches 10 mV/ms, is significantly more hyperpolarized in rats compared to mice (rat: -47.5±5.5 mV [SD], n=66; mouse: -40.0±5.3 mV [SD], n=118; P=4.9 × 10^−15^, unpaired, two-tailed t-test) (Fig. 3G).

**Figure 3.**
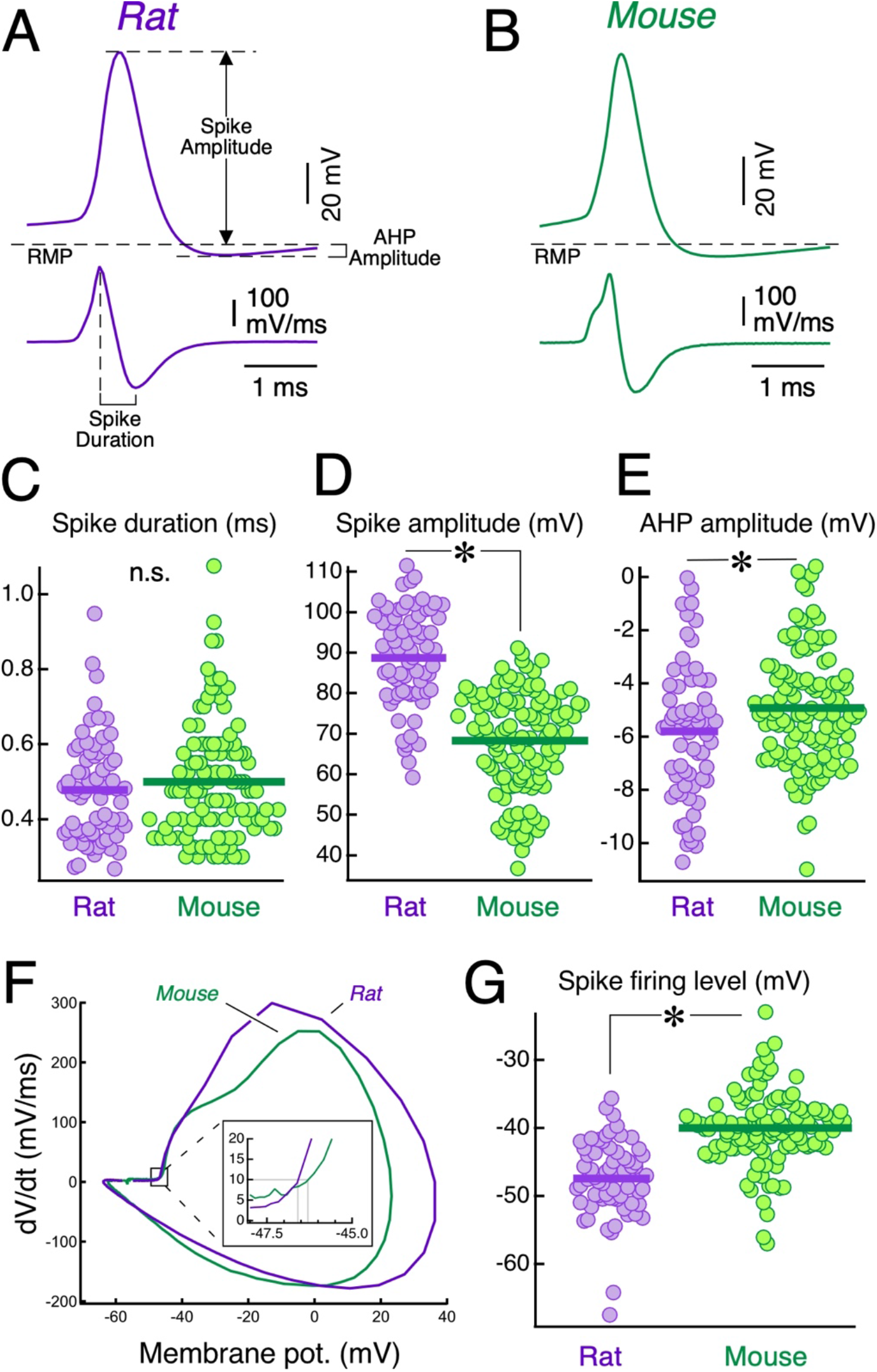
Spike characteristics of MesV neurons. *A* and *B*: Representative action potentials from rat and mouse respectively (top), with their corresponding first time derivative (bottom). Spike and spike after-hyperpolarization (AHP) amplitudes were measured from resting membrane potential level (RMP) to their peaks, whereas spike duration was measured as the time difference between maximum and minimum values of time derivative. *C*: Spike duration measured in the population of recorded neurons in rats and mice (rat: 0.48±0.14 ms [SD], n=69; mouse: 0.50±0.15 ms [SD], n=117; P=0.316, unpaired, two-tailed *t*-test). *D*: Spike amplitude measured in the population of recorded neurons in rats and mice (rat: 88.7±11.9 mV [SD], n=70; mouse: 68.2±12.3 mV [SD], n=118; P=1.29 × 10^−21^, unpaired, two-tailed *t*-test). *E*: AHP amplitude in rats and mice (rat: -5.8 ±2.5 mV [SD], n=68; mouse: -4.9±2.1 mV [SD], n=118; P=0.019, unpaired, two-tailed *t*-test). *F*: Phase plots of the first time derivative of the membrane potential (dV/dt) against the instantaneous membrane potential, for the traces depicted in *A* and *B*, shown superimposed. *Inset*: larger scale of the boxed area in the phase plots. The firing level is indicated for each trace (gray lines). *G*: Plot of the firing level in rats and mice (rat: -47.5±5.5 mV [SD], n=66; mouse: -40.0±5.3 mV [SD], n=118; P=4.9 × 10^−15^, unpaired, two-tailed t-test). Horizontal bars in *C, D, E* and *G* represent population averages.

The above result raises the possibility that the higher rate of postsynaptic recruitment in rats might be a consequence of presynaptic spikes larger in amplitude. However, their faster time course, corresponding to larger high-frequency components in the frequency domain, might result in more dramatic attenuation due to the filter properties of electrical synaptic transmission. Consistent with a previous study (Curti *et al*., 2012), frequency-transfer characteristics determined by ZAP protocols and FFT analysis (see Methods), revealed that despite some degree of frequency preference, electrical transmission between MesV neurons from both species essentially obeys a low-pass filter, with an apparent cutoff frequency of 51 Hz and 40 Hz in rats and mice respectively (Fig. 4A). According to these results, predicting the relative efficacy of transmission in these two species is not straightforward, as spikes of larger amplitude might result in larger postsynaptic coupling potentials, whereas attenuation of high frequency components due to low-pass filter properties is expected to have the opposite effect.

**Figure 4.**
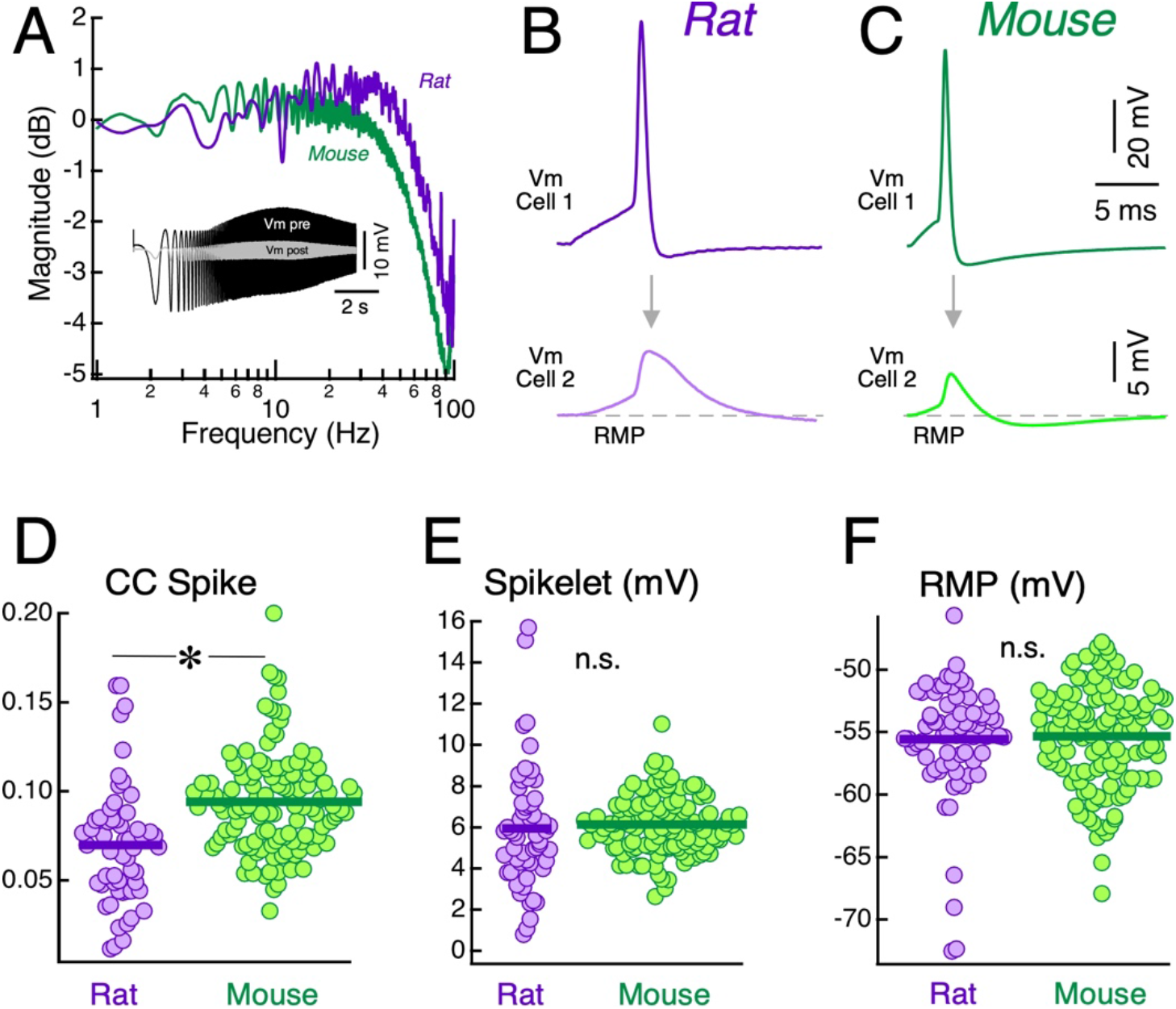
Spike transmission properties between coupled MesV neurons. *A*: Magnitude of frequency-transfer function between MesV neurons from rats and mice. Each curve represents the average of 14 directions in rats and 34 directions in mice. *Inset*: representative membrane responses to ZAP protocols of the pre- and postsynaptic neurons, from which the FFT were determined. *B* and *C*: Spike transmission in rat and mouse respectively, showing the presynaptic spike (above, Vm Cell1) evoked by a short depolarizing current pulse and corresponding coupling potentials in the postsynaptic neurons (below, Vm Cell 2). *D*: Spike related coupling coefficient (CC Spike) for the population of assessed directions in rats and mice calculated from recordings like those depicted in *B* and *C* (rat: 0.070±0.034 [SD], n=57; mouse: 0.094±0.028 [SD], n=122; P=7.66 × 10^−6^, unpaired, two-tailed *t*-test). *E*: Spike evoked coupling potential (Spikelet) amplitudes recorded in rats and mice (rat: 5.95±2.91 mV [SD], n=57; mouse: 6.13±1.41 mV [SD], n=122; P=0.645, unpaired, two-tailed *t*-test). *F*: Resting membrane potential (RMP) measured in the population of recorded neurons in rats and mice (rat: -55.6±4.6 mV [SD], n=70; mouse: -55.3±4.0 mV [SD], n=128; P=0.696, unpaired, two-tailed *t*-test). Horizontal bars in *D, E* and *F* represent population averages.

To directly assess the efficacy of spike transmission in these two species, the spike-related CC (CC Spike) for the population of electrically coupled MesV neurons was compared. For this, the CC Spike was determined as the ratio between the postsynaptic coupling potential (spikelet) amplitude, and the amplitude of the presynaptic spike from recordings like those depicted in Figure 4B and C. Noteworthy, this analysis revealed that the CC Spike from mice is almost 30% larger than that from rats, averaging 0.094±0.028 [SD] (n=122) and 0.070±0.034 [SD] (n=57) respectively (P=7.66 × 10^−6^, unpaired, two-tailed *t*-test) (Fig. 4D), indicating that spike transmission is more efficient in mice compared to rats. This difference most probably results from the fact that spikes from rats seem to present larger high-frequency components in comparison to mice (see above). However, despite this difference in CC spike, spikelet’s amplitudes from these two populations show no statistical difference, as they averaged 5.95±2.91 mV [SD] (n=57) in rats, and 6.13±1.41 mV [SD] (n=122) in mice (P=0.645, unpaired, two-tailed *t*-test) (Fig. 4E). These results indicate that higher efficiency of spike transmission in mice is compensated by the lower amplitude of presynaptic spikes, resulting in postsynaptic coupling potentials of similar amplitude. Moreover, it is worth mentioning that the resting membrane potential of MesV neurons from these two species do not show statistical difference (rat: -55.6±4.6 mV [SD], n=70; mouse: -55.3±4.0 mV [SD], n=128; P=0.696, unpaired, two-tailed *t*-test) (Fig. 4F).

Preceding results show that in both species presynaptic spikes evoke postsynaptic coupling potentials similar in amplitude, which arise from similar resting membrane potential level, suggesting that difference in postsynaptic recruitment should result from dissimilarities in membrane excitability. To test this possibility, a detailed characterization of firing properties was performed by using experimental protocols consisting of a series of depolarizing current pulses of increasing intensity (50 to 600 pA; Fig. 5A and B). From these experiments, plots of the number of spikes as a function of the injected current intensity were constructed (Fig. 5C). The average behavior of the population of rats and mice can be seen in Fig. 5D. This graph shows that rat MesV neurons respond with more spikes in almost the entire range of current intensity tested. Consistently, the slope of linear regression to spikes vs. current relationships in rats is significantly higher than in mice, averaging 0.011±0.015 spikes/pA [SD] (range: 0.0004 – 0.0664, n=70) and 0.003±0.008 spikes/pA [SD] (range: 0.0004 – 0.0637, n=99) respectively (P=7.19 × 10^−5^, unpaired, two-tailed *t*-test) (Fig. 5E), whereas the threshold current for the population of MesV neurons is significantly lower in rats than in mice, averaging 205.7±104.4 pA [SD] (n=70), and 396.2±181.4 pA [SD] (n=128) respectively (P=1.12 × 10^−16^, unpaired, two-tailed *t*-test) (Fig. 5F). These results clearly indicate that rat MesV neurons present a higher excitability, which is consistent with their lower firing level and suggest that it might underlie the higher efficacy in postsynaptic recruitment.

**Figure 5.**
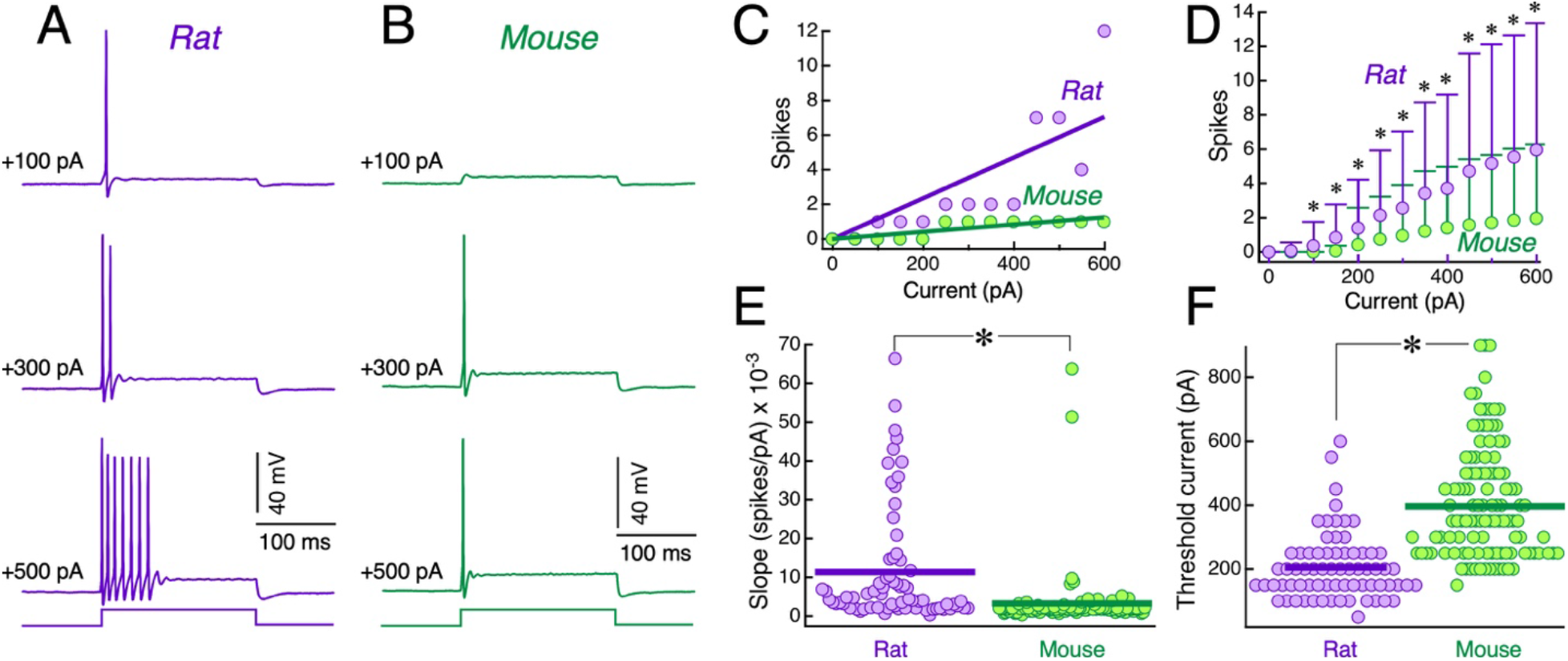
Firing properties of MesV neurons. *A* and *B*: Representative responses of a MesV neuron to intracellular depolarizing current pulses of increasing magnitude from rat and mouse respectively. *C*: Plot of the number of spikes evoked by current pulses of 200 ms in duration as a function of the injected current intensity for the same neurons depicted in *A* and *B*. Linear regression fits are shown superimposed to each data set *D*: Plot of the mean number of spikes evoked by current pulses (200 ms duration) as a function of the injected current intensity, for the population of recorded MesV neurons from rats (n=68, purple symbols) and mice (n=99, green symbols). Error bars represent SD. For each current intensity, statistical difference between rats and mice was evaluated by using unpaired, two-tailed *t*-test (0 pA: P=N/A; 50 pA: P=N/A; 100 pA: P= 0.0023; 150 pA: P=4.8×10^−5^; 200 pA: P= 0.0302; 250 pA: P= 0.0160; 300 pA: P= 0.0190; 350 pA: P= 0.0058; 400 pA: P= 0.0052; 450 pA: P= 0.0017; 500 pA: P= 0.0006; 550 pA: P= 0.0004; 600 pA: P= 0.0002). *E*: Slope values of linear regression fitted to spikes vs. current relationships like those depicted in *C*, for rats (purple symbols) and mice (green symbols) (rat: 0.011±0.015 spikes/pA [SD], range: 0.0004 – 0.0664, n=70; mouse: 0.003±0.008 spikes/pA [SD], range: 0.0004 – 0.0637, n=99; P=7.19 × 10^−5^, unpaired, two-tailed *t*-test). *F*: Plot of the threshold current for the population of recorded MesV neurons from rats and mice (rat: 205.7±104.4 pA [SD], n=70; mouse: 396.2±181.4 pA [SD], n=128; P=1.12 × 10^−16^, unpaired, two-tailed *t*-test). Horizontal bars in *E* and *F* represent population averages.

To gain insights into the membrane mechanism responsible for such difference in cellular excitability between rats and mice, voltage clamp experiments were designed in order to characterize the main subthreshold K+ currents (IA and ID), whose involvement in membrane resonance, oscillations, and bursting has been well established in these neurons (Del Negro & Chandler, 1997; Wu *et al*., 2001; Enomoto *et al*., 2006; Saito *et al*., 2006; Hsiao *et al*., 2009; Yang *et al*., 2009). Accordingly, IA and ID of MesV neurons from both species were characterized following standard protocols (see Methods). Figure 6A shows representative results from rats and mice, in which IA was recorded in voltage clamp, during protocols consisting in a series of step commands from 0 to 70 mV in steps of 5 mV and starting from a holding potential of ∼ -70 mV. Membrane currents obtained following this procedure showed rapid activation and an inactivation process with a kinetics typical of IA (Fig. 6B) (Storm, 1990; Mitterdorfer & Bean, 2002). From membrane current recordings, conductance values were calculated in order to construct activation curves which were fitted to a Boltzmann function (Fig. 6C). From these fits, maximum conductance (gmax), half activation voltage (Vhalf) and slope at Vhalf were obtained for the population of recorded neurons. This analysis revealed that IA from rats and mice present similar characteristics and cannot account for differences in membrane excitability between these species. In fact, gmax normalized by the cell’s capacitance to avoid bias by difference in cell size, averaged 1.40±0.61 nS/pF [SD] (n=17) in rats and 1.42±0.38 nS/pF [SD] (n=14) in mice (P=0.944, unpaired, two-tailed *t*-test) (Fig. 6D, left), Vhalf averaged -27.9±4.4 mV [SD] (n=17) in rats and -28.9±5.2 mV [SD] (n=14) in mice (P=0.571, unpaired, two-tailed *t*-test) (Fig. 6D, middle), whereas slope averaged 9.51±1.71 mV [SD] (n=17) in rats and 9.55±2.61 mV [SD] (n=14) in mice (P=0.969, unpaired, two-tailed *t*-test) (Fig. 6D, right).

**Figure 6.**
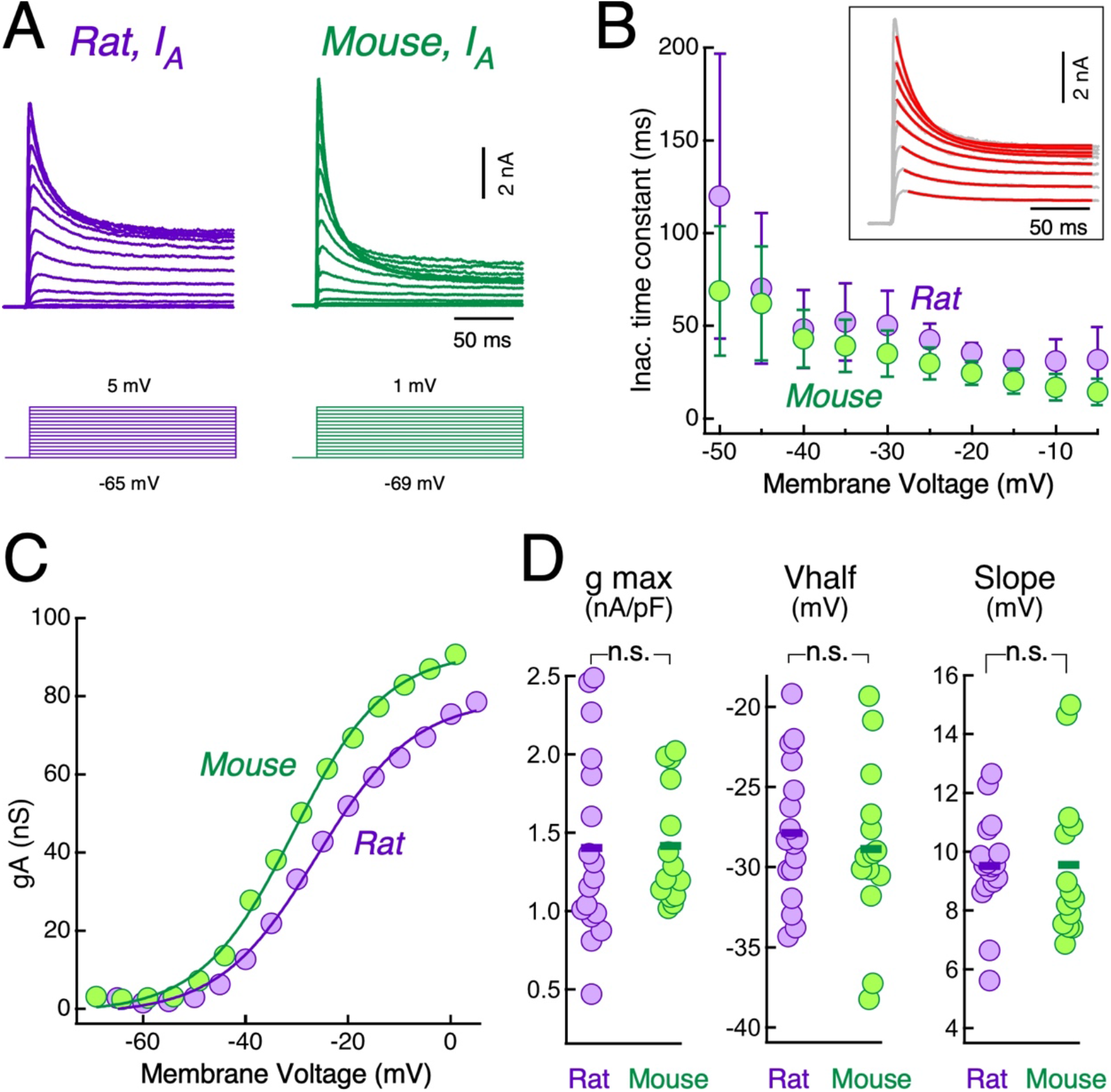
Characterization of the A-type current of MesV neurons. *A*: Representative traces of IA current recorded in a MesV neuron from rat (left) and mouse (right). Voltage commands employed are illustrated below current traces. *B*: Plot of inactivation mean time constant as a function of voltage for the population of recorded MesV neurons from rats (n=13, purple symbols) and mice (n=11, green symbols). Error bars represent SD. *Inset*: fittings to single exponential functions (red traces) to the falling phase of membrane currents during voltage commands (grey traces) in order to estimate the inactivation time constants. *C*: Activation curves constructed from traces shown in *A*. Fits to a Boltzmann function (continuous trace) are superimposed to the experimental data (round symbols). *D*: Plots of the IA maximal conductance normalized by the cell’s capacitance (gmax, left) (rat: 1.40±0.61 nS/pF [SD], n=17; mouse: 1.42±0.38 nS/pF [SD], n=14; P=0.944, unpaired, two-tailed *t*-test), half activation voltage (Vhalf, middle) (rat: -27.9±4.4 mV [SD], n=17; mouse: -28.9±5.2 mV [SD], n=14; P=0.571, unpaired, two-tailed *t*-test) and slope values at Vhalf (Slope, right) (rat: 9.51±1.71 mV [SD], n=17; mouse: 9.55±2.61 mV [SD], n=14; P=0.969, unpaired, two-tailed *t*-test) obtained from fits to Boltzmann function, for the population of recorded neurons. Horizontal bars represent population averages.

Representative recordings of ID from rats and mice are shown in Figure 7A. This conductance is characterized by its rapid activation and a slow voltage-dependent inactivation kinetics (Fig. 7B). ID inactivation is considerable slower than that of the IA, confirming that they were successfully separated by the experimental protocols. Strikingly, in contrast to the IA, ID gmax is 63% higher in MesV neurons from mice compared to rats, averaging 2.26±0.90 nS/pF [SD] (n=13) and 1.39±0.56 nS/pF [SD] (n=20) respectively (P=0.0057, unpaired, two-tailed *t*-test) (Fig. 7C and 7D, left). Whereas Vhalf (rat: -39.2±3.8 mV [SD], n=20; mouse: -36.3±4.2 mV [SD], n=13; P=0.056, unpaired, two-tailed *t*-test) (Fig. 7D, middle) and slope at Vhalf (rat: 6.92±2.42 mV [SD], n=20; mouse: 8.54±2.70 mV [SD], n=13; P=0.092, unpaired, two-tailed *t*-test) (Fig. 7D, right), did not exhibit statistical difference. Comparison of average activation curves for the population of recorded neurons, confirms the higher functional expression of ID in mice (Fig. 7E). Moreover, its incomplete inactivation determines a large window current of about 50-60% of gmax at membrane voltages between -40 and -30 mV, and about 10% close to the RMP in both species (Fig. 7F and G), supporting the notion that ID contributes to set the Rin and the RMP. This is consistent with a more depolarized firing level and lower excitability of mice MesV neurons (Bekkers & Delaney, 2001; Guan *et al*., 2007; Higgs & Spain, 2011; Ordemann *et al*., 2019). On the other hand, the INap, whose involvement has also been established as a critical determinant of MesV neurons’ excitability in both species (Wu *et al*., 2001; Enomoto *et al*., 2006, 2007), showed no significant difference between rats and mice. Indeed, INap gmax averaged 0.096±0.025 nS/pF [SD], (n=10) and 0.12±0.075 nS/pF [SD] (n=8) in rats and mice respectively (P=0.445, unpaired, two-tailed *t*-test). The Vhalf averaged -45.9±4.5 mV [SD] (n=10) and -43.2±8.2 mV [SD] (n=8) in rats and mice respectively (P=0.412, unpaired, two-tailed *t*-test), whereas the slope at Vhalf averaged 5.1±1.0 mV [SD] (n=10) and 6.0±2.4 mV [SD] (n=8) in rats and mice respectively (P=0.356, unpaired, two-tailed *t*-test) (Supp. Fig. 1).

**Figure 7.**
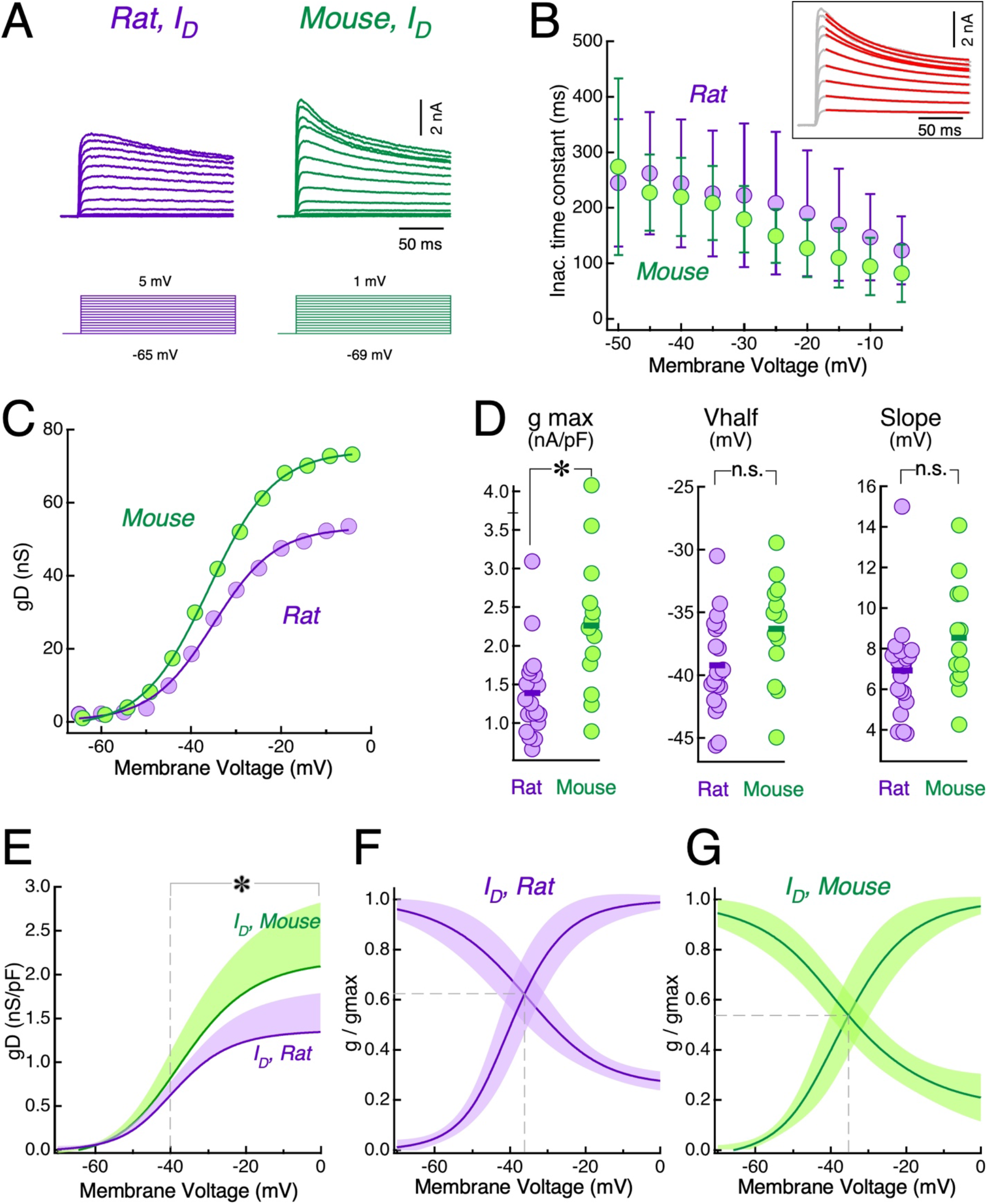
Characterization of the D-type current of MesV neurons. *A*: Representative traces of ID current recorded in a MesV neuron from rat (left) and mouse (right). Voltage commands employed are illustrated below current traces. *B*: Plot of inactivation mean time constant as a function of voltage for the population of recorded MesV neurons from rats (n=13, purple symbols) and mice (n=11, green symbols). Error bars represent SD. *Inset*: fittings to single exponential functions (red traces) to the falling phase of membrane currents during voltage commands (grey traces) in order to estimate the inactivation time constants. *C*: Activation curves constructed from traces shown in *A*, fits to Boltzmann function (continuous traces) are superimposed to the experimental data (round symbols). *D*: Plots of the ID maximal conductance (gmax, left) (rat: 1.39±0.56 nS/pF [SD], n=20; mouse: 2.26±0.90 nS/pF [SD], n=13; P=0.0057, unpaired, two-tailed *t*-test), half activation voltage (Vhalf, middle) (rat: -39.2±3.8 mV [SD], n=20; mouse: -36.3±4.2 mV [SD], n=13; P=0.056, unpaired, two-tailed *t*-test) and slope values at Vhalf (Slope, right) (rat: 6.92±2.42 mV [SD], n=20; mouse: 8.54±2.70 mV [SD], n=13; P=0.092, unpaired, two-tailed *t*-test) obtained from fits to Boltzmann function, for the population of recorded neurons. Horizontal bars represent population averages. *E:* Plot shows the average activation curves of the ID for the population of recorded neurons in rats (purple) and mice (green). Conductance values were normalized by the cell’s capacitance. Shaded area represents SD. *F* and *G*: Activation and inactivation curves of the ID current from rats and mice respectively. Each curve represents the average of fits to Boltzmann function for the population of recorded neurons normalized by its maximum values. Shaded area represents SD. Vertical dashed line indicates the intersection between activation and inactivation curves corresponding to the maximum “window” current.

Taken together, the preceding results strongly suggest that the higher level of ID expression in MesV neurons from mice underlies their lower excitability in relation to rat MesV neurons. To confirm this interpretation, the involvement of ID in regulating the cellular excitability was directly assessed by a pharmacological approach. Figure 8A and B show results from mice MesV neurons in which the addition of 4-AP (30 μM) to block ID, resulted in a marked increase in firing for the entire range of injected current intensity. In fact, these neurons that typically respond to depolarizing current pulses with one or two spikes in control conditions, in the presence of 4-AP respond with robust repetitive responses. Consistently, the slope of linear regressions of spikes vs. current relationships displayed a significant increase from 0.0015±0.0004 spikes/pA [SD] in control, to 0.012±0.010 spikes/pA [SD] after 4-AP addition (n=19, P=0.00019, paired, two-tailed *t*-test) (Fig. 8B and C), whereas the threshold current was significantly reduced from 389.5±117.4 pA [SD] in control, to 78.9±41.9 pA [SD] in the presence of 4-AP (n=19, P=1.89×10^−10^, paired, two-tailed *t*-test) (Fig. 8D). These results clearly indicate that the ID plays a critical role regulating membrane excitability, and its expression level contributes to defining the electrophysiological phenotype of MesV neurons in a species-specific fashion. To test whether the control of membrane excitability exerted by ID also impact on the efficacy of postsynaptic recruitment, the effect of 4-AP (30 μM) was assessed on the ability of presynaptic action potentials to drive spiking in postsynaptic coupled neurons in mice. Results from these experiments are illustrated in Figure 8E, in which spiking in the presynaptic neuron fails to activate the coupled neuron in control conditions (left), whereas after addition of 4-AP (30 μM) there is a dramatic increase in firing of the postsynaptic cell (right). In fact, while in control conditions recruitment of the postsynaptic neuron was not observed in any of the tested directions, after blockade of the ID, recruitment occurred in 11 out of 19 directions, close to the proportion observed in rats in control conditions (see Fig.1C). Consistently, the average number of postsynaptic spikes in relation to presynaptic spikes varied from 0.0±0.0 % [SD] in control to 19.7±30.4 % [SD] in 4-AP (n=19; P=0.011, paired, two-tailed *t*-test) (Fig. 8E, inset). These results clearly indicate a critical role of the ID in controlling the efficacy of postsynaptic recruitment, and hence in determining the operation mode of electrical synapses between MesV neurons.

**Figure 8.**
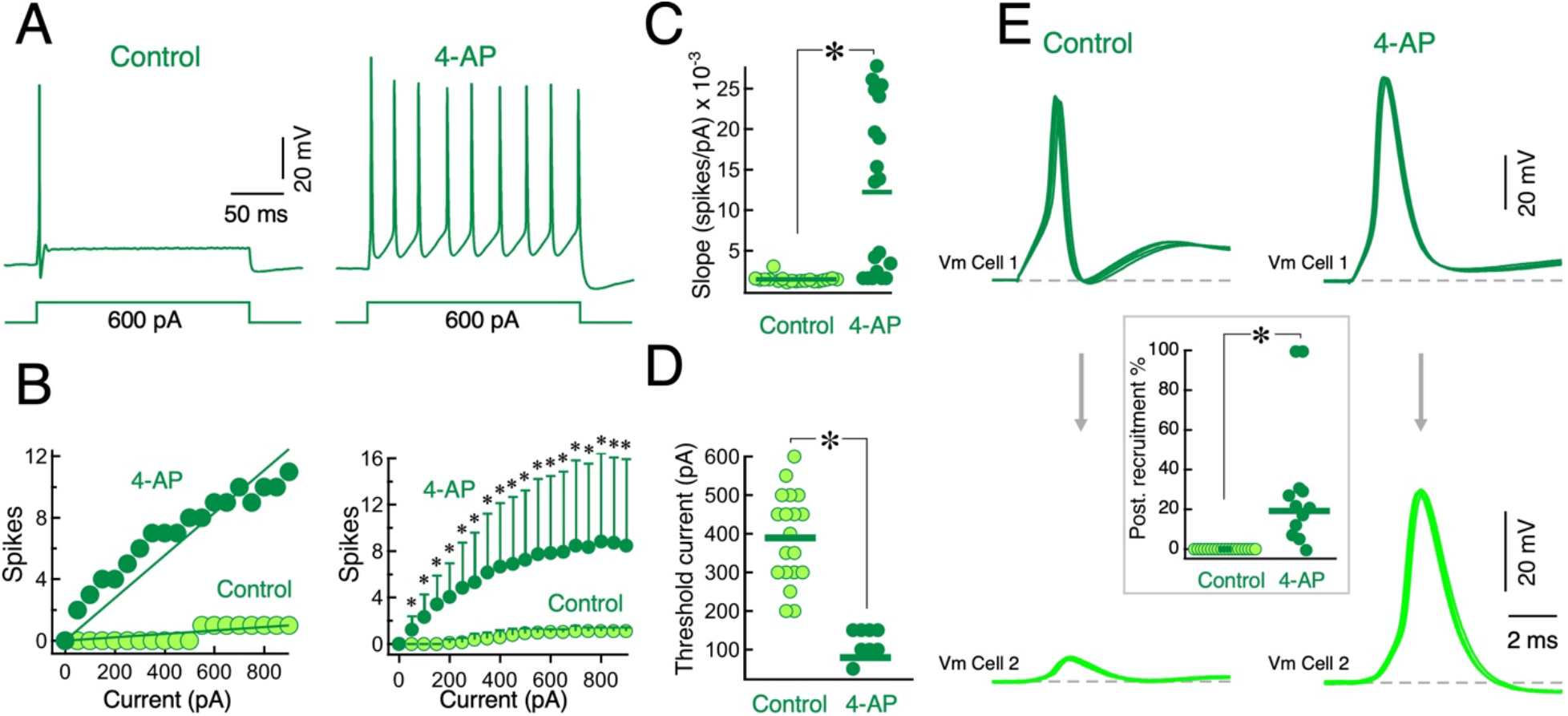
Blocking ID increases excitability and efficacy of postsynaptic recruitment in mice. *A*: Typical response of a MesV neuron from mouse to a suprathreshold depolarizing pulse in control conditions (left), and after the addition of 4-AP (30 μM) to the bath solution (right), in order to block ID current. Below voltage traces are depicted the injected current pulses. *B*: Plots of the number of spikes evoked by current pulses of 200 ms in duration as a function of the injected current intensity for the same neuron depicted in *A* (left) and for the population of recorded MesV neurons in mice, in control conditions (light green symbols) and in the presence of 4-AP (30 μM) (dark green symbols). Error bars represent SD. For each current intensity, statistical difference between rats and mice was evaluated by using paired, two-tailed *t*-test (0 pA: P=N/A; 50 pA: P= 0.0002; 100 pA: P= 7.7×10^−5^; 150 pA: P= 1.6×10^−5^; 200 pA: P= 1.1×10^−5^; 250 pA: P= 3.5×10^−5^; 300 pA: P= 5.5×10^−5^; 350 pA: P= 7.7×10^−5^; 400 pA: P= 8.5×10^−5^; 450 pA: P= 0.0002; 500 pA: P= 0.0002; 550 pA: P= 0.0003; 600 pA: P= 0.0003; 650 pA: P= 0.0004; 700 pA: P= 0.0004; 750 pA: P= 0.0004; 800 pA: P= 0.0004; 850 pA: P= 0.0003; 900 pA: P= 0.0005). *C*: Slope values of linear regression fitted to spikes vs. current relationships as depicted in part *B* at left, in control conditions (light green symbols) and in the presence of 4-AP (30 μM) (dark green symbols) (control: 0.0015±0.0004 spikes/pA [SD]; 4-AP: 0.012±0.010 spikes/pA [SD], n=19; P=0.00019, paired, two-tailed *t*-test). *D*: Threshold current of MesV neurons from mice in control conditions (light green symbols) and in the presence of 4-AP (30 μM) (dark green symbols) (control: 389.5±117.4 pA [SD]; 4-AP: 78.9±41.9 pA [SD], n=19; P=1.89×10^−10^, paired, two-tailed *t*-test). *E*: Representative results obtained in a pair of electrically coupled MesV neurons in which the presynaptic neuron was activated with a short depolarizing current pulse (Vm Cell 1), while recording the corresponding membrane voltage response in the postsynaptic coupled neuron (Vm Cell 2), in control conditions (left) and in the presence of 4-AP (30 μM) (right). 50 – 100 single traces are shown superimposed in order to evaluate the incidence of spiking in the postsynaptic coupled neuron. *Inset*: Plot of the efficacy of postsynaptic recruitment quantified as the fraction of postsynaptic spikes in relation to presynaptic spikes expressed as percentage in control conditions (light green symbols) and in the presence of 4-AP (30 μM) (dark green symbols) (control: 0.0±0.0 % [SD]; 4-AP: 19.7±30.4 % [SD], n=19; P=0.011, paired, two-tailed *t*-test).

Further insights into the mechanism by which such control is exerted, was obtained by comparing the spikelets from these two species. Typical examples from rats and mice are illustrated in Figure 9A and B respectively. Despite similar peak amplitudes, their waveforms present important differences. Rats spikelets display a falling phase characterized by large variability and delayed time to peak, resulting in significant longer duration (Fig. 9C). Indeed, spikelet half-amplitude duration averaged 4.6±1.8 ms [SD] (n=22) and 2.4±0.7 ms [SD] (n=106) in rats and mice respectively (P=2.03×10^−5^, unpaired, two-tailed *t*-test) (Fig. 9D). This protracted duration is consistent with the participation of the INap, whose activation promotes spiking of the postsynaptic coupled neurons in rats (Curti & Pereda, 2004; Curti *et al*., 2012). In contrast, in mice MesV neurons, which on average express 63% more ID, the INap is strongly antagonized by the swift activation of this outward current, curtailing the spikelets duration and therefore reducing the efficacy of postsynaptic recruitment. To confirm this hypothesis, artificial spikelets were generated in single MesV neurons from mice, by injecting current waveforms corresponding to postjunctional currents measured in voltage clamp during paired recordings. For this, pairs of electrically coupled MesV neurons were simultaneously recorded, one cell in current clamp and the other in voltage clamp (Fig. 9E, inset). While spiking was induced in the current-clamped cell by means of short depolarizing current pulses (Fig. 9E, upper and middle traces), the resultant membrane current (postjunctional current) was recorded in the postsynaptic voltage-clamped neuron (Fig. 9E, lower trace). During independent recordings from single cells, the opposite (sign reversed) of the postjunctional recorded current was injected as a current clamp command, and its intensity scaled according to the neuron’s Rin in order to obtain a spikelet of typical amplitude (Fig. 9F, inset). Such artificial spikelets, that were indistinguishable from real ones (spikelet amplitude: 6.13±1.41 mV [SD], n=122; artificial spikelet amplitude: 6.63±0.95 [SD], n=17; P=0.073, unpaired, two-tailed *t*-test), were generated in control conditions and in the presence of 4-AP (30 μM), in order to evaluate the contribution of the ID to its waveform (Fig. 9F). In those cases, in which 4-AP application depolarized the neuron’s RMP, it was corrected to near pre-application levels by injecting DC current. This approach was chosen instead of assessing spike transmission in current clamp during paired recordings because ID also contributes to spike repolarization (Fig. 8E) (Mitterdorfer & Bean, 2002), and changes in presynaptic spike waveform would also affect spikelet duration. Confirming our hypothesis, half-amplitude duration of artificial spikelets was significantly increased after ID blockade, averaging 2.0±0.3 ms [SD] and 5.3±5.3 ms [SD] in control and in the presence of 4-AP (30 μM) respectively (P=0.023, paired, two-tailed *t*-test; n=17). Moreover, recruitment by artificial spikelets increased from 0.0±0.0 % [SD] in control to 9.1±16.4 % [SD] in the presence of 4-AP (30 μM) (P=0.037, paired, two-tailed *t*-test; n=17). Altogether, these results suggest that the density of ID determines coupling potentials duration and postsynaptic recruitment in electrically coupled MesV neurons.

**Figure 9.**
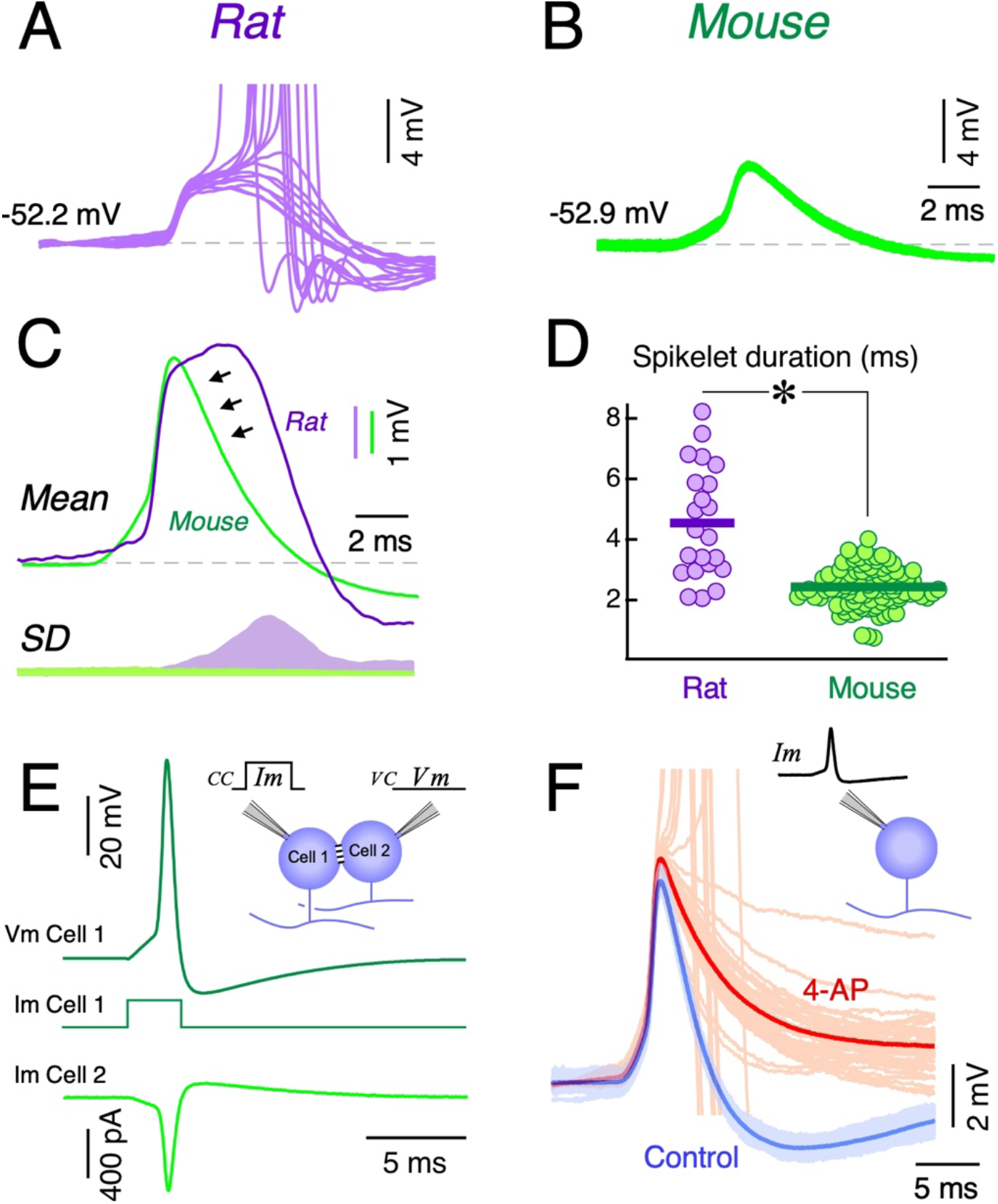
ID current expression determines spikelet time course. *A* and *B*: Typical spike-evoked postsynaptic coupling potentials in MesV neurons from rats and mice respectively. Successive responses are shown superimposed during repetitive activation of the presynaptic neuron. *C*: Superimposed traces illustrate the averaged postsynaptic potentials (Mean, above) and its standard deviation (SD, below) corresponding to the recordings in A and B. Traces with spikes were not included. *D*: Spikelet half-amplitude duration for rats (4.6±1.8 ms [SD], n=22) and mice (2.4±0.7 ms [SD], n=106) (P=2.03×10^−5^, unpaired, two-tailed *t*-test). *E*: Paired recordings from electrically coupled mouse MesV neurons, one cell in current clamp (Cell 1) and the other in voltage clamp (Cell 2). The current-clamped cell was activated (upper trace, Vm Cell 1) by means of short depolarizing current pulses (middle trace, Im Cell 1), and the resultant membrane current (junctional current) was recorded in the postsynaptic voltage-clamped neuron (lower trace, Im Cell 2). The inset illustrates the experimental paradigm. *F*: Artificial spikelets in a single mouse MesV neuron evoked by injecting a current waveform corresponding to the junctional current (inset) in control conditions (light blue: single traces, dark blue: average trace) and in the presence of 4-AP (30 μM) (light red: single traces, dark red: average trace from single traces without spikes).

## DISCUSSION

The comparative study of the MesV nucleus from rats and mice exposed a remarkable species-specific difference in the operation mode of electrical synapses, revealing the role of the intrinsic electrophysiological properties in shaping the behavior of circuits of coupled neurons. In the present study, a naturally occurring difference in the efficacy of postsynaptic recruitment between electrically coupled MesV neurons was used to uncover the critical role of subthreshold K+ currents. We showed that, while in rats the spiking of one MesV neuron activates its coupled partner in ∼50% of the cases, this rarely occurs between mice MesV neurons, in which spiking results in a subthreshold postsynaptic response in approximately 98% of the tested pairs. Noteworthy, the population of connected MesV neurons from both species bear similar strength of electrical coupling, as a result of comparable magnitude of its determinants (junctional conductance and neuronal input resistance). Thus, despite spike-evoked postjunctional coupling potentials (spikelets) from both species being of similar amplitude and arising from a comparable resting membrane potential level, postsynaptic recruitment is considerably more efficient in rats than in mice. This striking difference is imposed by the differential expression of the D-type K+ current, whose density is significantly higher in MesV neurons from mice compared to rats. Consistently, previous reports suggested that firing properties of MesV neurons result from the relative expression of the INap and the subthreshold K+ currents (Wu *et al*., 2001, 2005; Hsiao *et al*., 2009). Regarding to this, MesV neurons have been shown to express several types of Kv1 subunits, whose pharmacological profile and voltage dependency, matches those of the ID characterized in the present study. In fact, α-DTX, considered as a specific blocker of Kv1 mediated currents, show an almost complete overlap in the blocking effect with 4-AP in the low micromolar range (50 μM) (Saito *et al*., 2006; Hsiao *et al*., 2009).

Previous experimental and theoretical work has already shown the relevance of active membrane properties levering the activity of networks of coupled neurons, particularly through the action of boosting mechanisms like the INap (Mann-Metzer & Yarom, 1999; Pfeuty *et al*., 2003; Curti & Pereda, 2004; Dugué *et al*., 2009; Curti *et al*., 2012). The synergic operation of these mechanisms endows neural circuits with the ability to generate synchronic and rhythmic patterns of activity with potential functional relevance for both, physiological and pathophysiological processes (Draguhn *et al*., 1998; Perez Velazquez & Carlen, 2000; Hormuzdi *et al*., 2001; Mylvaganam *et al*., 2014). Moreover, other circuit operations supported by electrical synapses like coincidence detection are also critically shaped by the intrinsic excitability of neurons. Indeed, the ability of coupled neurons to discriminate between synchronic and temporally uncorrelated inputs, is strongly regulated by the hyperpolarization activated cationic current (H-current) (Davoine & Curti, 2019). In spite of this background, direct experimental evidence for the role of K+ currents in the context of electrical synaptic transmission was lacking. Here, we show that the expression level of the D-type K+ current, a depolarization-activated subthreshold conductance, whose activation kinetics ranges the time scale of physiologically relevant signals like action potentials, gates the transfer of spikes between coupled neurons. In fact, the swift activation of this outward current at near the resting membrane potential opposes depolarizations, curtailing the duration of spike-evoked coupling potentials and regulating membrane excitability.

Previous work has shown that the synergistic interaction between electrical coupling and the intrinsic neuronal properties promotes the strong synchronic activation of MesV neurons (Curti *et al*., 2012). Interestingly, while the efficacy of postsynaptic recruitment in rats is high, facilitating the spread of excitation among MesV neurons, the incidence of electrical coupling is relatively low (∼23% of apposed pairs are electrically coupled in rats compared to ∼63% in mice) (Curti *et al*., 2012). Thus, cellular excitability and electrical coupling at the MesV nucleus of these species are inversely related, suggesting that the expression of these mechanisms are reciprocally regulated in a sort of homeostatic relationship. In this regard, regulations at the network level that reciprocally operates on the expression of mechanisms underlying cellular excitability and interneuronal connectivity supported by gap junctions, might be critical to ensure network function stability, as was proposed in networks of neurons interconnected by chemical contacts (Marder & Goaillard, 2006). Indeed, widespread coupling between highly excitable neurons might lead to hypersynchrony and aberrant spiking across neuronal ensembles, characteristic of diseases like epilepsy (Perez Velazquez & Carlen, 2000).

MesV neurons are primary sensory afferents that originate in spindles of jaw-closing muscles (masseter) and mechanoreceptors of periodontal ligaments (Kolta *et al*., 1990; Westberg *et al*., 2000). In turn, similarly to their spinal cord counterparts, MesV neurons establish monosynaptic excitatory connections with motoneurons controlling the same muscles, contributing to the organization of orofacial behaviors (Luo & Li, 1991; Grimwood *et al*., 1992; Luo *et al*., 2001; Stanek *et al*., 2014). By virtue of electrical synapses, activation of one afferent might lead to the activation of a coupled partner, increasing the number of active afferents that respond coordinately to a sensory input, thus supporting lateral excitation as was shown in many sensory systems (El Manira *et al*., 1993; Herberholz *et al*., 2002; Rela & Szczupak, 2004). In this way, this phenomenon, whose expression is under control of the ID, enhances or amplifies the influence of sensory input on jaw-closing motoneurons.

Finally, these findings illustrate how related species display different cellular and circuital strategies to solve the same biological problem, in this case the control of orofacial behaviors. Similar diversity in homolog circuits from rats and mice has also been shown in hypothalamic neurons controlling pituitary prolactin secretion, whose networks display contrasting behaviors. However, in contrast to our findings, such diversity arises from interspecific differences in electrical coupling, while neurons from both species present comparable membrane properties (Stagkourakis *et al*., 2018).

## Acknowledgements

We thank A. Pereda, G. Budelli, F. Trigo and V. Abudara for critical discussions and comments.

## Grants

This work was supported by Agencia Nacional de Investigación e Innovación (ANII), Uruguay (FCE_1_2021_1_166745), Programa de Desarrollo de las Ciencias Básicas (PEDECIBA) and Comisión Académica de Posgrado of Universidad de la República.

## Disclosure

The authors declare no conflict of interests.

---

**Supplemental Figure 1.**
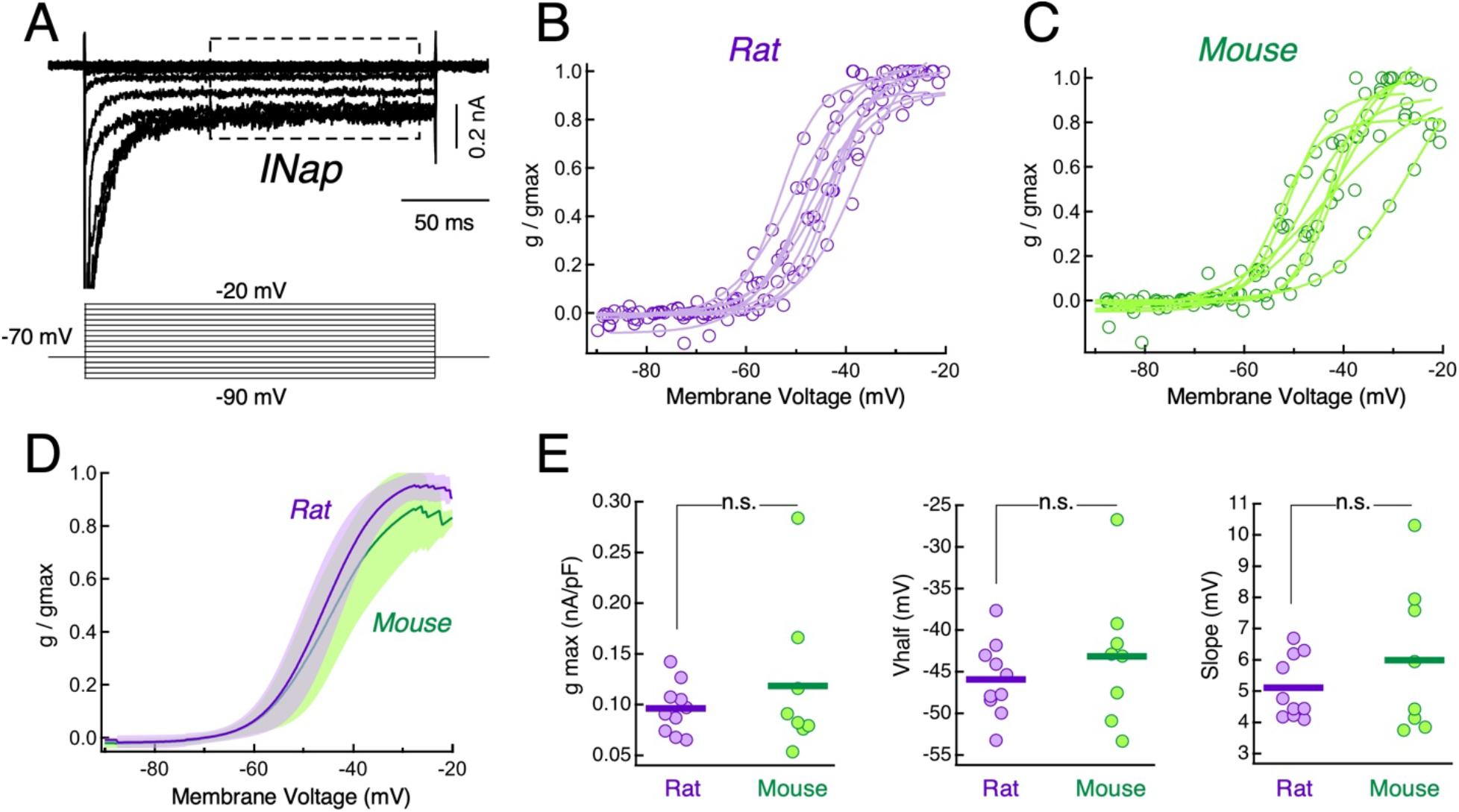
Characterization of the persistent Na^+^ current of MesV neurons. *A*: Representative traces of the INap recorded in a MesV neuron (above) and the voltage commands employed (below). *B* and *C*: Activation curves from the population of recorded MesV neurons from rats and mice respectively. Experimental values (round symbols) and fits to a Boltzmann function (continuous traces) are shown superimposed. Conductance values determined from the TTX-sensitive and non-inactivating component of membrane currents (boxed area in *A*), were normalized to it maximum values. *D*: Activation curves of the INap from rats (purple) and mice (green). Each curve represents the average of fits to Boltzmann function for the population of recorded neurons normalized by its maximum values. Shaded area represents SD. *E*: Plots of the INap maximal conductance, normalized by the cell’s capacitance (gmax, left) (rat: 0.096±0.025 nS/pF [SD], n=10; mouse: 0.12±0.075 nS/pF [SD], n=8; P=0.445, unpaired, two-tailed *t*-test), half activation voltage (Vhalf, middle) (rat: -45.9±4.5 mV [SD], n=10; mouse: -43.2±8.2 mV [SD], n=8; P=0.412, unpaired, two-tailed *t*-test) and slope values at Vhalf (Slope, right) (rat: 5.1±1.0 mV [SD], n=10; mouse: 6.0±2.4 mV [SD], n=8; P=0.356, unpaired, two-tailed *t*-test) obtained from fits to Boltzmann function, for the population of recorded neurons. Horizontal bars represent population averages.

## Notes

### Competing Interest Statement

The authors have declared no competing interest.

